# Characterization of the cluster MabR prophages of *Mycobacterium abscessus* and *Mycobacterium chelonae*

**DOI:** 10.1101/2022.04.27.489733

**Authors:** Jacob Cote, Colin Welch, Madeline Kimble, Dakota Archambault, John Curtis Ross, Hector Orellana, Katelyn Amero, Claire Bourett, Andre Daigle, Keith W. Hutchison, Sally D. Molloy

**Affiliations:** Department of Molecular and Biomedical Sciences, University of Maine, Orono, Maine; The Honors College, University of Maine, Orono, Maine

**Keywords:** prophage, Mycobacterium, bacteriophage, genome

## Abstract

*Mycobacterium abscessus* is an emerging pathogen of concern in cystic fibrosis and immunocompromised patients and is considered one of the most drug-resistant mycobacteria. The majority of clinical *M. abscessus* isolates carry one or more prophages that are hypothesized to contribute to virulence and bacterial fitness. The prophage McProf was identified in the genome of the Bergey strain of *M. chelonae*, and is distinct from previously described prophages of *M. abscessus*. The McProf genome increases intrinsic antibiotic resistance of *M. chelonae* and drives expression of the intrinsic antibiotic resistance gene, *whi*B7, when superinfected by a second phage. The prevalence of McProf-like genomes was determined in sequenced mycobacterial genomes. Related prophage genomes were identified in the genomes of 25 clinical isolates of *M. abscessus* and assigned to the novel cluster, MabR. The MabR genomes share less than 10% gene content with previously described prophages; however, share features typical of prophages, including polymorphic toxin immunity (PT-Imm) systems.

## Introduction

Prophages are viral genomes integrated into bacterial genomes and they contribute to the genetic diversity and virulence of many bacterial pathogens (Brüssow et al. 2004; Costa et al. 2018; Figueroa-Bossi et al. 2001; Fortier 2018; Fortier and Sekulovic 2013; Wang and Wood 2016). Clinically important non-tuberculosis mycobacteria (NTM), such as *Mycobacterium abscessus*, often cause drug-resistant infections and continue to be a significant public health burden (Nasiri et al. 2017). The majority of clinical NTM carry prophage genomes that are enriched in genes that potentially promote bacterial fitness and virulence (Dedrick et al. 2021; Glickman et al. 2020).

The prophages of *M. abscessus* are highly diverse and distinct from the mycobacteriophage genomes in the Actinobacteriophage database of phagesdb.org (Dedrick et al. 2021; Russell and Hatfull 2016). Dedrick et al. (2021) identified 122 prophage sequences in 82 clinical isolates of *M. abscessus* of which 67 were unique (*Dedrick et al. 2021*). These were sorted into 17 Mab clusters (MabA – MabQ) based on shared gene content (>35% shared genes) (Dedrick et al. 2021). Many of the prophages encode toxin/antitoxin (TA) and polymorphic toxin/immunity (PT-Imm) systems that are hypothesized to contribute to virulence (Dedrick et al. 2021; Zhang et al. 2012). We recently described a novel prophage genome, named McProf, in the genome of *M. chelonae* (*M. chelonae* CCUG 47445 coordinates 1,521,426 – 1,589,648) that shares only 10% gene content with the Dedrick et al. prophages, but encodes numerous genes expressed during lysogeny, including a PT-Imm system (Cushman et al. 2021). McProf contributes to the intrinsic drug resistance of *M. chelonae* and increases expression of the conserved mycobacterial regulator of intrinsic antibiotic resistance genes, *whi*B7, when superinfected by a second mycobacteriophage. Understanding the prevalence of this novel prophage genome and its relationship with known prophage genomes will be important for a better understanding of the role of prophage genomes in mycobacterial fitness and virulence.

In this study, prophage genomes related to McProf were identified in 26 published genomes of mycobacteria. Gene content was compared with prophage genomes described by Dedrick et al. (2021) and sorted into a novel cluster, MabR (Dedrick et al. 2021). Here we report the genomes of 5 unique cluster MabR genomes, including 4 *M. abscessus* prophages and the original *M. chelonae* prophage McProf.

## Materials and methods

### Identification and Extraction of Prophage from Mycobacterial genomes

Prophage sequences similar to McProf were identified using the PhagesDB BLASTn tool to search *M. abscessus* genomes within the PATRIC database (Altschul et al. 1990; Russell and Hatfull 2016; Wattam et al. 2014). Highly scoring sequences were analyzed using PHASTER to determine the putative coordinates of prophage genomes within bacterial genome sequences (Arndt et al. 2016). Precise coordinates were determined after manual inspection of prophage genomes and identification of repeat sequences that flank the prophage genome and represent the common core of *attL/attR* sites. Each prophage sequence was extracted with the identified attachment sites defining the genome ends. Prophages were named according to the strain in which they reside, i.e., prophiXXXX01-1, with suffixes used to denote multiple prophages in the same genome as described by Dedrick et al. 2021 (Dedrick et al. 2021).

### Prophage genome annotation and comparative genomics

Prophage genes were predicted using Glimmer and GeneMark within DNA Master (http://cobamide2.bio.pitt.edu/) and PECAAN (Borodovsky et al. 2003; Delcher et al. 1999; Rinehart et al. 2016). The start site for each gene was determined through manual inspection. Gene functions were predicted using the web-based tools HHpred and NCBI BLASTp (Altschul et al. 1990; Söding et al. 2005). Dot plots were constructed using gepard using default settings (Krumsiek et al. 2007). The prophage network phylogeny is based on shared gene content and was created in SplitsTree (Huson 1998). Genome maps were created using Phamerator and the “Actino_Mab_Draft” database, version 19 (Cresawn et al. 2011). Integration sites were predicted by comparing flanking bacterial sequence in each prophage genome to that of *M. abscessus* ATCC 19977. Specific integration locations were determined by probing the previous integration region with the *attL* sequence for each prophage. Alignments with 100% sequence identity were considered to be core *attB* sites.

## Results

### Identification of cluster MabR prophage

In order to identify prophage sequences related to the *M. chelonae* prophage McProf, we searched the NCBI database using BLASTN and identified a prophage sequence in the *Mycobacterium phlei* strain NCTC8151 (accession number LR134347) with 100% nucleotide identity to the McProf genome. To search for McProf-like sequences in *M. abscessus* genomes, we probed the PATRIC database with the McProf genome sequence using the BLASTN feature within phagesdb.org (Altschul et al. 1990; Russell and Hatfull 2016; Wattam et al. 2014). We identified 25 *M. abscessus* clinical strains carrying prophage sequences with high sequence identity to the McProf genome (Table 1). All of the *M. abscessus* strains were isolated from the respiratory system of diseased individuals, and the vast majority of the *M. abscessus* strains were isolated in the United Kingdom (76%) (Table 1). The remaining 24% of strains were isolated in the United States (16%) and Australia (8%).

**Table 1.**
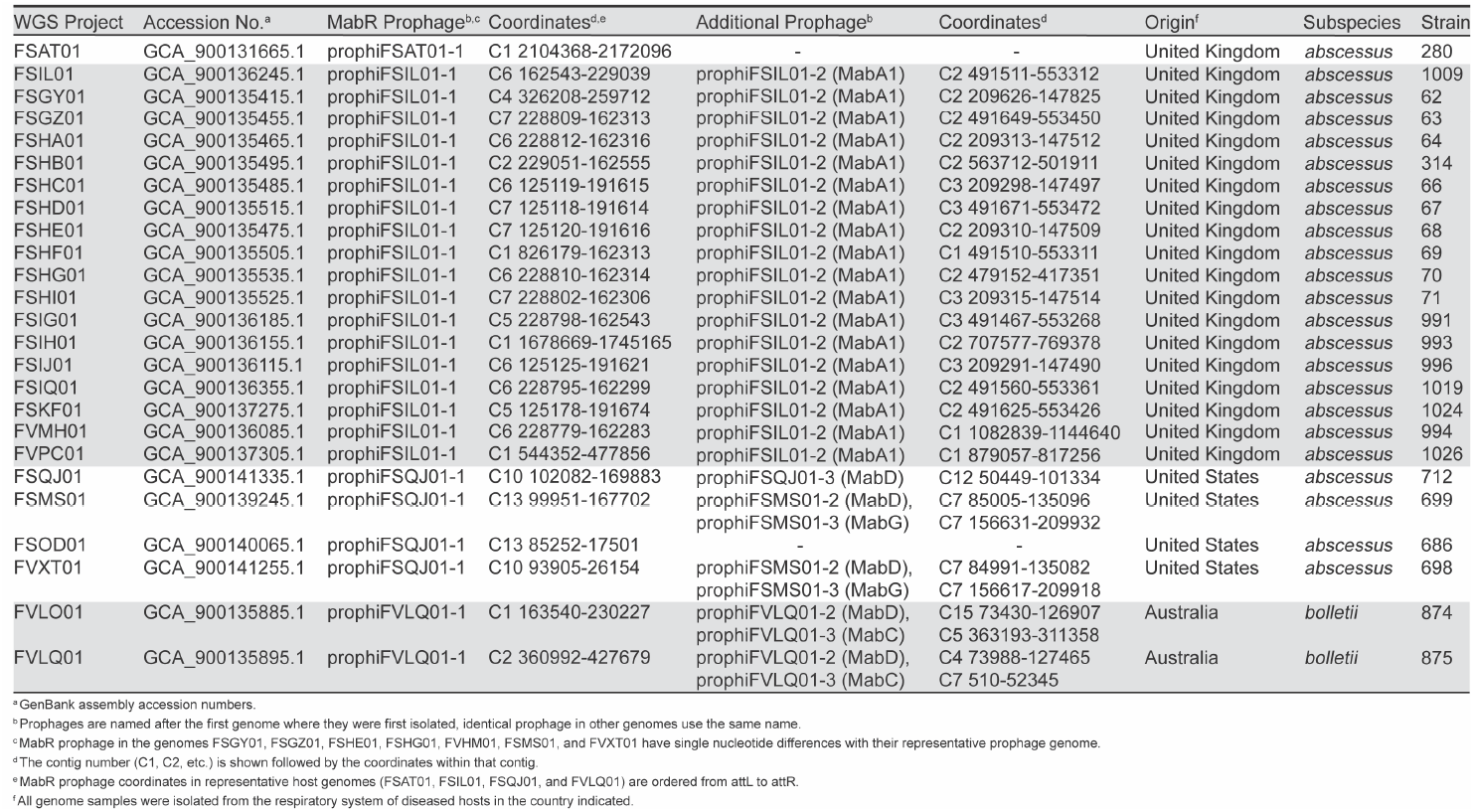
*M. abscessus* bacterial strains carrying cluster MabR prophage

Of the 25 identified McProf-like prophage sequences, only four prophage sequences were unique. These four prophage sequences were extracted from the bacterial sequences of the following *M. abscessus* strains: FSAT01, FSIL01, FSQJ01, and FVLQ01 (Table 2). The ends of the prophage genomes were determined by the left and right attachment sites flanking the prophage genomes (Table 2). Prophages were named by the strain they were extracted from and the number of prophages identified in the strain: prophi[strain]-# (Table 2). McProf and the four McProf-like prophage genomes: prophiFSAT01-1, prophiFSIL01-1, prophiFSQJ01-1, and prophiFVLQ01-1 share less than 10% genome content with the *M. abscessus* prophages described by Dedrick et al. (2021) and were assigned to a novel cluster, MabR (Figure 1A) (Dedrick et al. 2021). The MabR prophages overall have high nucleotide similarity to one another (Figure 1B).

**Table 2.**
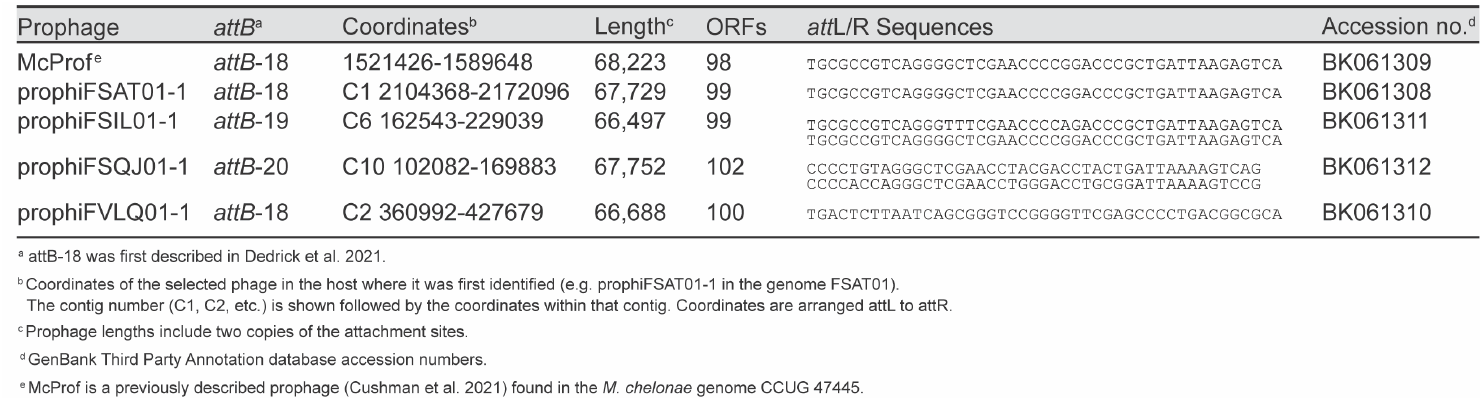
Genome characteristics of cluster MabR prophages and select cohabitating prophages

**Figure 1.**
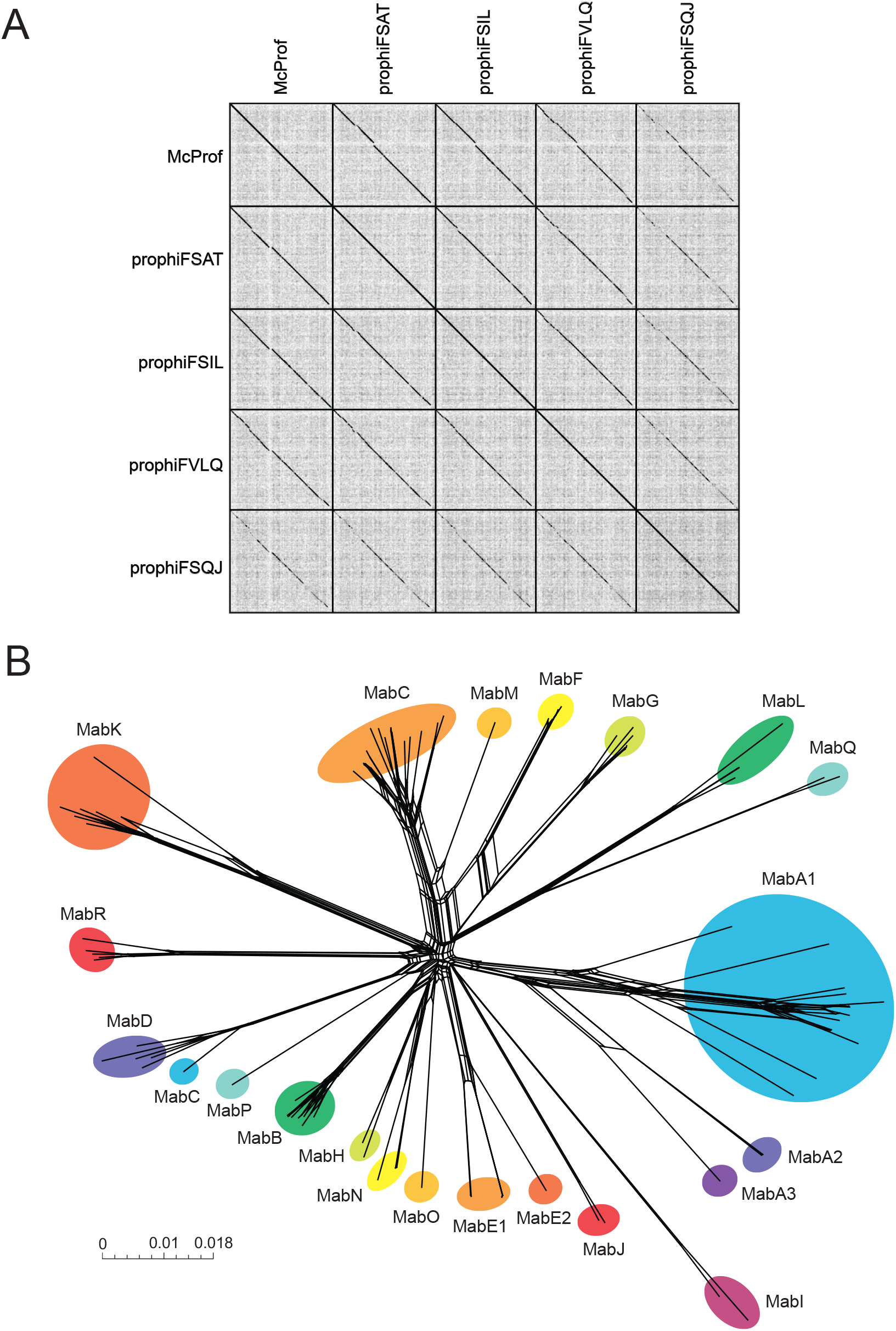
Diversity of MabR prophages. (A) Dotplot comparison of MabR prophages. (B) Phylogenetic network representation of cluster MabR prophages and *M. abscessus* prophages (Dedrick et al. 2021) based on shared gene content as described by Pope et al. 2015. Nodes represent individual prophage; circles represent prophage clusters. Scale marker indicates substitutions/site.

### Integration locations

The integration sites of MabR prophage were determined and compared to that of prophage described by Dedrick et al. (Dedrick et al. 2021). Most of the prophage genomes integrated into known *M. abscessus attB* sites, often in the 3’ end of tRNA genes, but several utilized novel integration sites (Table 3). Three prophage genomes, McProf, prophiFSAT01-1, and prophiFVLQ01-1, integrate into the 3’ end of a tRNA-Lys (*attB*-18) as described in Dedrick et al. (2021) (Figure 2). prophiFSIL01-1 integrates into the 3’ end of a tRNA-Lys (*attB*-19) and prophiFSQJ01-1 integrates into Mab_0771c (*attB*-20), a predicted major transport protein. *attB*-20 was the only cluster MabR integration site identified within a protein-coding sequence. The *attB* core sequences and coordinates for each identified integration site are listed in Table 3.

**Table 3.**
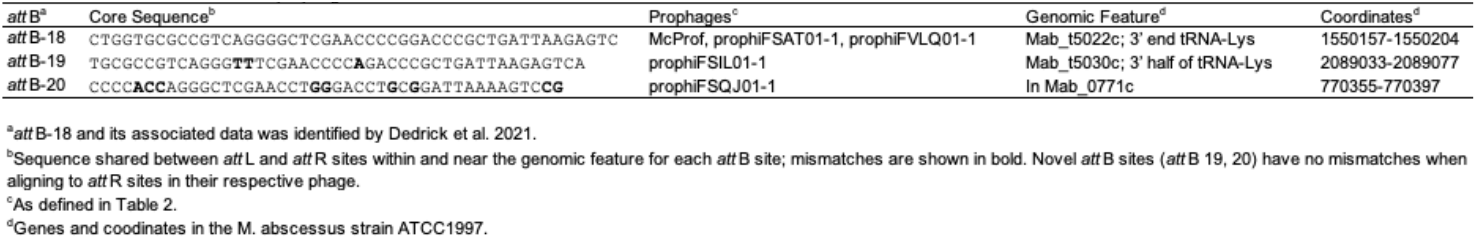
*att* B sites of cluster MabR prophage

**Figure 2.**
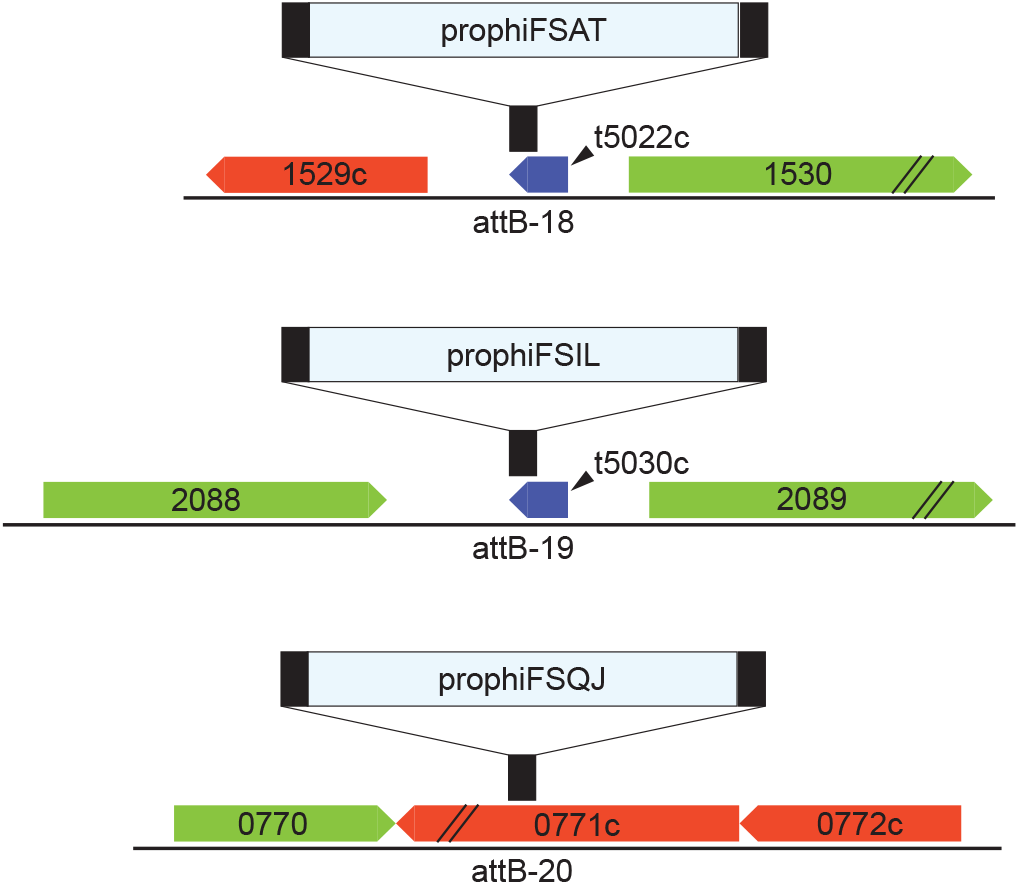
MabR prophage integration locations. The three integration schemes used by MabR prophage are shown as *attB* site locations (black bars) shown relative to *M. abscessus* ATCC 19977 genes for reference. Rightward and leftward transcribed genes are shown by green and red arrows, respectively, with their ATCC 19977 gene number. Both tRNAs (blue boxes) are transcribed in the leftward direction. Not shown are McProf and prophiFVLQ01-1 which utilize the *attB-18* site described by Dedrick et al. 2021.

### Genomic organization of cluster MabR genomes

MabR prophage have very similar genome architectures and areas of conserved gene content (Figure 3). The genomes are tightly packed, typical of mycobacteriophage genomes, containing 98-102 genes across approximately 67 kb. The integration and immunity cassettes are located immediately adjacent to the left attachment site (*attL*). All MabR genomes share a rightward transcribed tyrosine integrase (gp1), a gene of unknown function (gp2) and a leftward transcribed immunity repressor (gp3) (Figures 3 and 4). The immunity repressor is distinct from immunity repressors encoded by other Mab cluster prophages; however, is a homolog of the immunity repressors found in the genomes of five cluster K2 mycobacteriophage, DismalFunk, DismalStressor, Findely, Marcoliusprime and Milly. A Cro and excise gene (gp4 and 5) are divergently transcribed from the immunity repressor (Figures 3 and 4). The early lytic genes that follow show some diversity across the five MabR genomes, particularly in prophiFSQJ01-1. The structural, assembly, and lysis cassette genes are highly conserved across MabR genomes.

**Figure 3.**
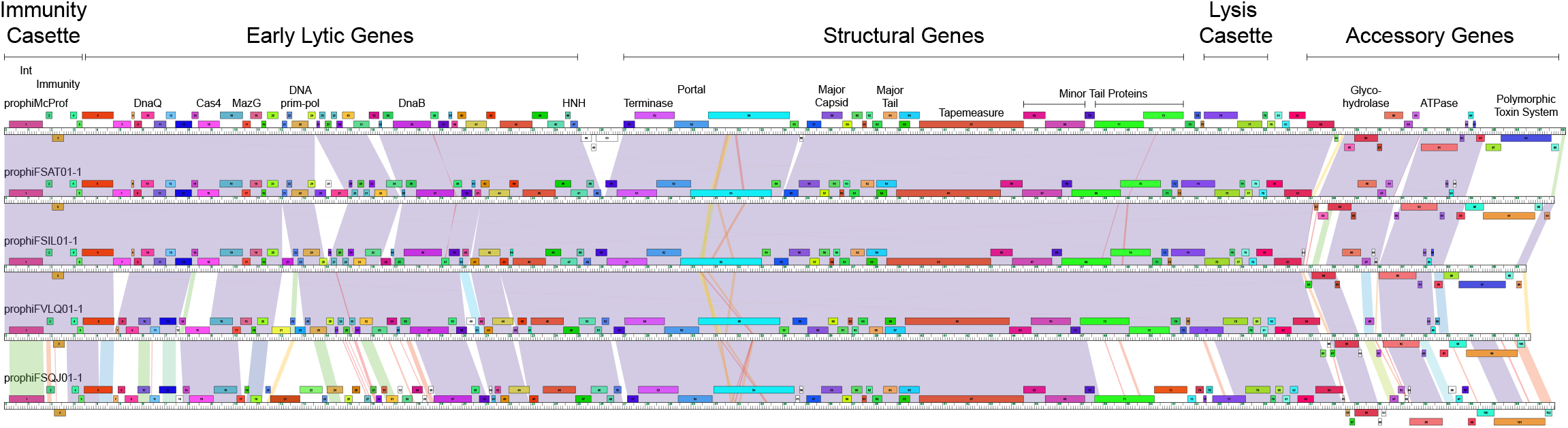
Organization of MabR genomes. MabR genomes are shown with pairwise nucleotide sequence similarity displayed by colors between genomes; purple is the most similar, red is the least similar above a BLASTN E threshold of 10^−5^. The ruler represents the coordinates of the genome. Forward and reverse transcribed genes are shown as boxes above and below the ruler, respectively. Maps were generated using Phamerator and the database, “Actino_Mab_Draft (version 20).” (Cresawn et al. 2011)

**Figure 4.**
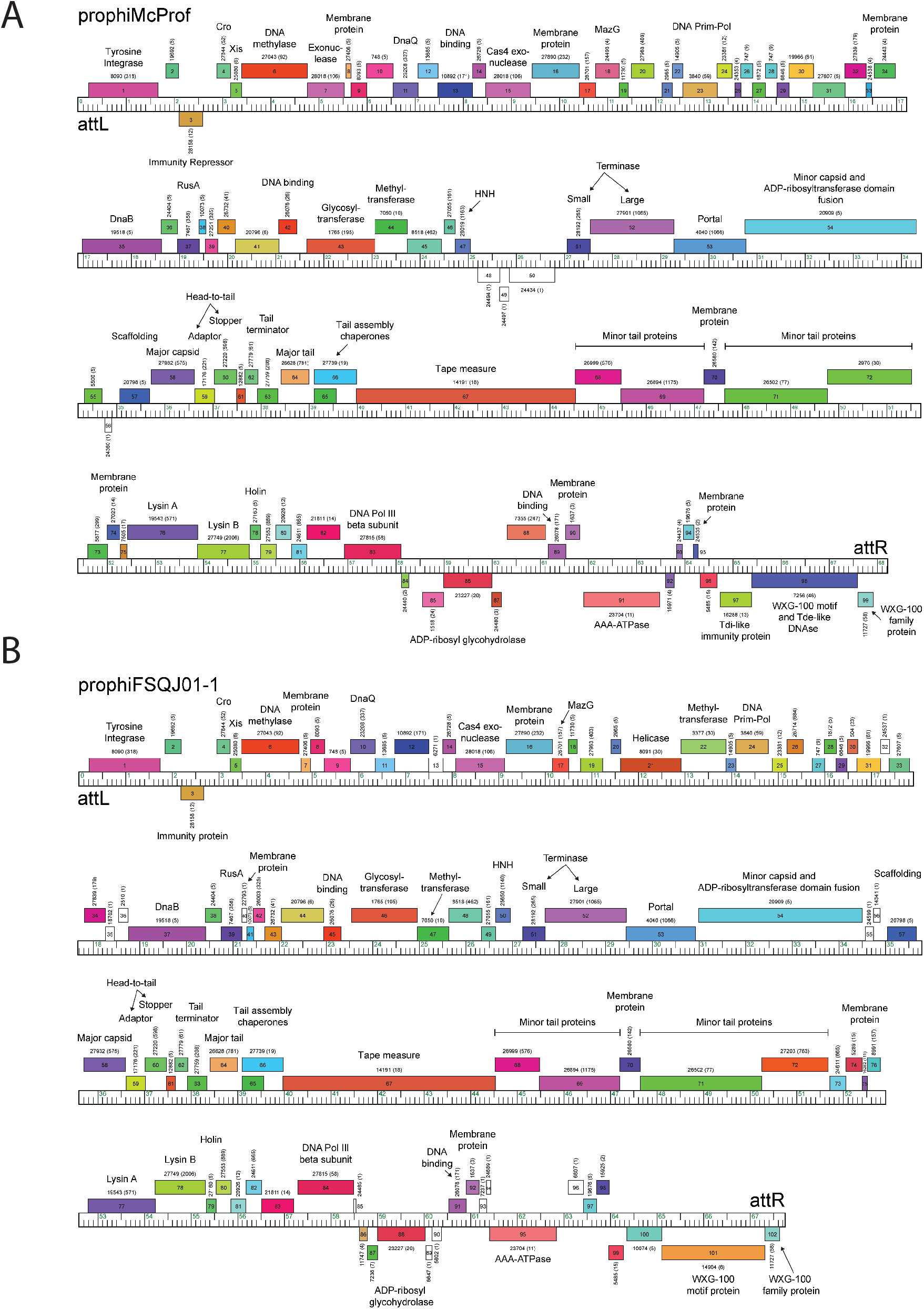
Genome organization of (A) prophiMcProf and (B) prophiFSQJ01-1. The ruler represents the coordinates of the genome. Forward and reverse transcribed genes are shown as boxes above and below the ruler, respectively. Genes are colored according to their assigned Phamily with the Phamily number shown above each gene with the number of Phamily members in parentheses. Genome maps were generated using Phamerator and the database, “Actino_Mab_Draft (version 20).” (Cresawn et al. 2011)

Between the lysis cassette and the right attachment site (*attR*) is a group of highly diverse genes that are most likely expressed during lysogeny (Figure 3) (Cushman et al. 2021; Dedrick et al. 2017). Some of the genes shared across all MabR genomes are unique to the cluster and include a DNA polymerase III sliding clamp, an ADP-ribosyl glycohydrolase, a helix-turn-helix DNA binding domain protein and an AAA-ATPase. Immediately adjacent to *attR*, all MabR prophage genomes encode a reverse transcribed polymorphic toxin immunity system (PT-Imm) that include an ESAT6-like WXG-100 protein, a polymorphic toxin and cognate immunity protein (Figures 3 and 5).

**Figure 5.**
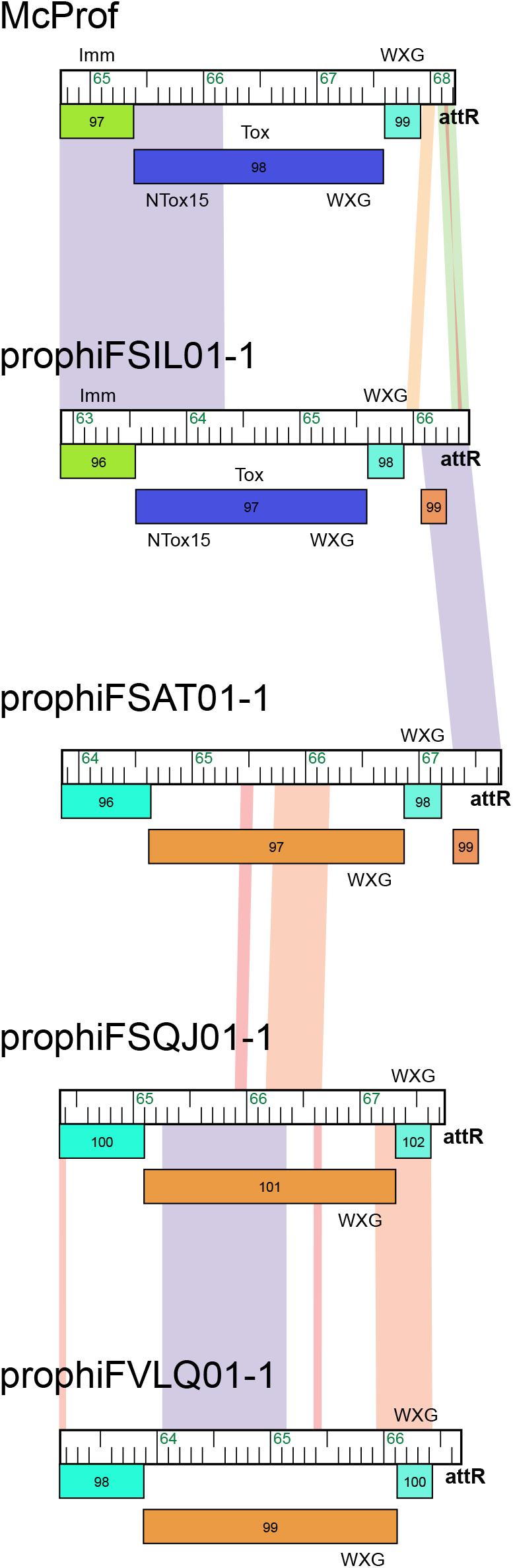
Organization of MabR polymorphic toxin-immunity systems. MabR genomes are aligned at their polymorphic toxin-immunity systems beginning at the 3’ end of the predicted immunity proteins. Genomes are displayed as described in Figure 2 but are ordered in such a way that genomes with the most similarity in this region are next to each other. Also shown are the motifs/domains found at the N- and C-termini of MabR polymorphic toxins. All predicted polymorphic toxins have a single WXG-100 motif at the N-terminus while the C-terminus is variable. Note that gp99 in prophiFSIL01-1 and prophiFSAT01-1 has no predicted function and is included to show the relationship of the polymorphic toxin systems to the genome ends.

### Polymorphic toxin systems

Dedrick et al. 2021 identified 21 distinct, highly modular, polymorphic toxin-immunity (PT-imm) systems across 50 *M. abscessus* prophage (Dedrick et al. 2021). These systems consist of a large polymorphic toxin (PT) and a cognate immunity protein to prevent self-toxicity and at least one ESAT6-like WXG-100 protein. The cluster MabR genomes contain one of two types of PT-Imm systems (Figure 3 and 5). The PT in the McProf and prophiFSIL01-1 genomes has an N-terminus WXG-100 motif and a C-terminus Tde-like DNAse toxin domain (Ntox15 PF15604) (Cushman et al. 2021; Ma et al. 2014). Downstream is the Tdi-like immunity protein with GAD-like and DUF1851 domains (Ma et al. 2014). This PT system is also found in the genome of prophiGD43A-5 (Figure 6). Although the three PT genes all carry the same Ntox15 domain, they share low sequence identity across the linker and WXG-100 domains. In the NCBI database this PT-Imm system is also found in the genomes of *Mycobacterium* phage phiT46-1 (Accession number NC_054432.1) and numerous mycobacterial species including *M. abscessus, M. goodie* and *M. salmoniphilum*.

**Figure 6.**
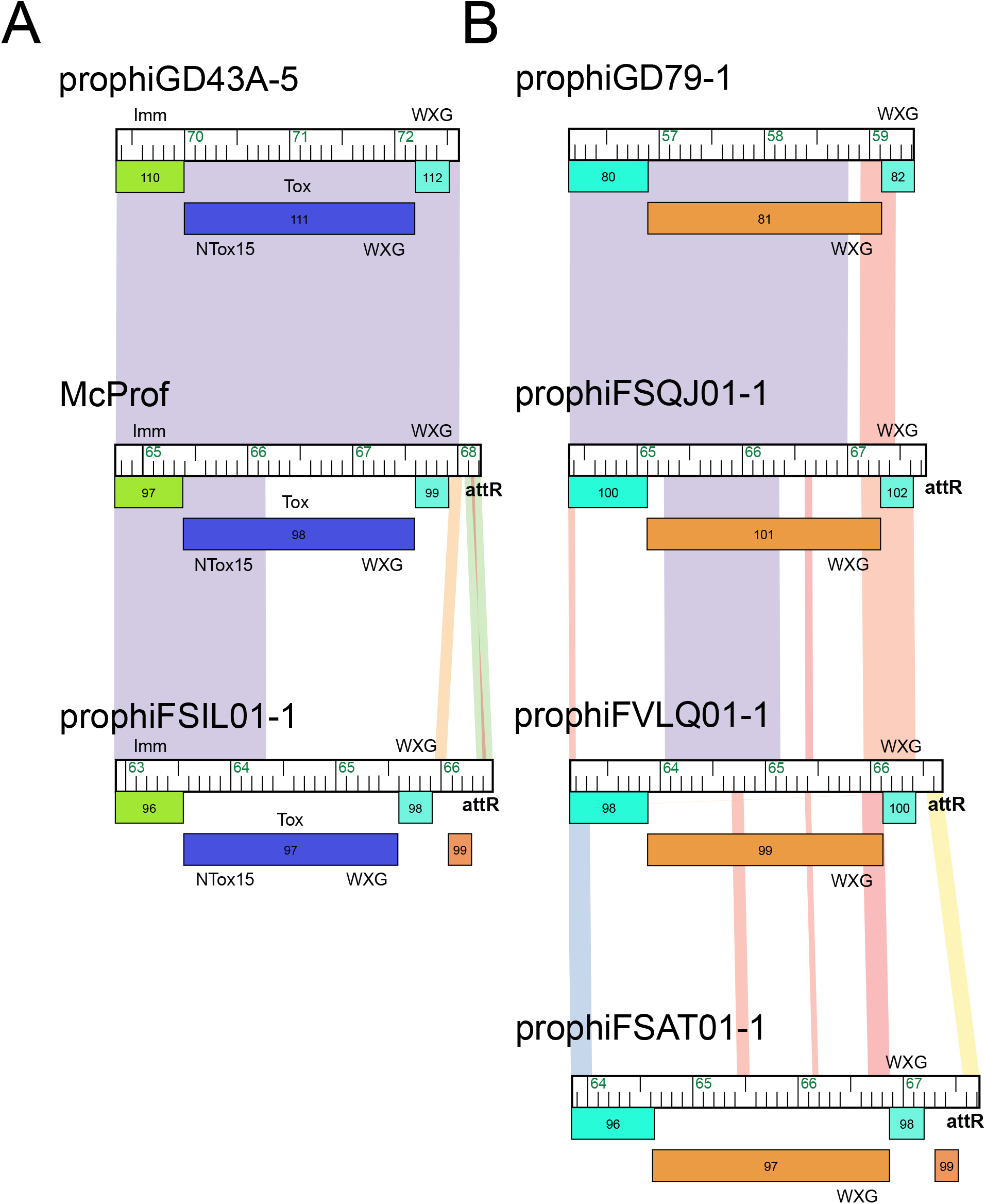
MabR polymorphic toxin-immunity systems found in non MabR prophage. Genomes are displayed as described in Figure 2 and Figure 5. prophiGD43-5 and prophiGD79-1 belong to clusters MabK and MabQ respectively.

The genomes of prophiFSAT01-1, prophiFSQJ01-1, and prophiFVLQ01-1 carry a gene cassette that is organized like a PT-Imm system and encodes an ESAT6-like WXG-100 protein (Figure 5). However, we were unable to predict toxin and immunity domains. The presumed PT gene has an N-terminus WXG-100 domain but lacks an identifiable toxin domain in the C-terminus. Likewise, the downstream gene lacks domains known to be associated with immunity, such as SUKH or Imm (Dedrick et al. 2021; Zhang et al. 2012). This second PT-Imm system is also found in the cluster MabQ genome, prophiGD79-1 (Figure 6) (Dedrick et al. 2021).

## Discussion

The majority of bacterial pathogens carry prophages that are known to contribute to bacterial virulence and fitness (Brüssow et al. 2004; Figueroa-Bossi et al. 2001; Wang and Wood 2016). Prophage introduce novel genes into bacterial genomes that can result in phenotypes that are more competitive in bacterial populations (Brüssow et al. 2004; Wang and Wood 2016). The prophage McProf is found in the Bergey strain of *M. chelonae* (ATCC 35752) and increases bacterial resistance to aminoglycosides (Cushman et al. 2021). Although the McProf genome is distinct from the *M. abscessus* prophages described by Dedrick et al. (2021) (Dedrick et al. 2021), it is clearly related to a novel subgroup of prophage genomes identified in the genomes of clinical *M. abscessus* isolates and therefore was assigned to the novel cluster, MabR.

The majority of the MabR prophages were identified in the genomes of *M. abscessus* isolates although a prophage genome that shared 100% nucleotide with McProf were identified in *M. phlei*. Of the 25 MabR genomes identified in *M. abscessus* strains, only four were unique and these were typically found in isolates with the same geographical origin (Table 1). Strains of the same geographic origin also typically carried identical co-habitating prophages, suggesting that the bacterial strains are highly related.

The MabR prophage genomes, although distinct in overall gene content, share a genome organization and some gene features that are typical of the prophages described by Dedrick et al (2021) (Dedrick et al. 2021). These include two types of PT-Imm systems that potentially contribute to mycobacterial fitness (Figures 5 and 6) (Zhang et al. 2012). The PT-Imm systems of McProf and prophiFSIL01-1 are similar to the PT-Imm system that plays a role in plant colonization in the Gram-negative plant pathogen, *Agrobacterium tumefaciens* (Ma et al. 2014). The polymorphic toxins share a C-terminal DNAse toxin domain (Ntox-15) but differ at the N-terminal domain which contains a WXG100 domain needed for interacting with Type VII secretion systems in mycobacteria versus the PAAR domain needed for Type VI secretion systems in *Agrobacterium* (Ma et al. 2014).

It’s not clear yet if the PT-Imm systems of the MabR prophage are important for bacterial fitness but it is known that the presence of the McProf genomes increases *M. chelonae* resistance to aminoglycosides relative to a non-lysogen strain (Cushman et al. 2021). The addition of a second prophage, cluster G phage BPs, to this strain further increased aminoglycoside resistance and increased expression of mycobacterial genes in the *whi*B7 regulon, including *whi*B7 (Cushman et al. 2021; Sampson et al. 2009). This large change in *whi*B7 expression and aminoglycoside resistance is driven by the presence of the McProf genome as it is not observed in strains carrying the BPs prophage alone. There are 16 genes expressed from the McProf genome during lysogeny of *M. chelonae* that potentially contribute to altered *whi*B7 expression and increased aminoglycoside expression (Cushman et al. 2021). Many of these genes are common across the MabR genomes including the McProf PT-Imm cassette (gp97 – 99), gp91 and 92, and gp85 and 86 (Figure 3). A better understanding of the function and role these genes potentially play in mycobacterial fitness will improve our overall understanding of how prophage contribute to mycobacterial virulence.

## Data availability

The bacterial genome coordinates of the MabR prophage genomes and the bacterial genome accession numbers are presented in Table 1. The genome sequences and annotations of prophages McProf (Accession No. BK061309), prophiFSAT01-1 (Accession No. BK061308), prophiFSIL01-1 (Accession No. BK061311), prophiFVLQ01-1 (Accession No. BK061310), and prophiFSQJ01-1 (Accession No. BK061312), are available through NCBI GenBank.

## Acknowledgements

We would like to thank Dr. Graham Hatfull, Christian Gauthier and Steven Cresawn for their support in phamerating prophage genomes and creating splits tree diagrams.

## Supplemental Data

**Figure SI** Genome organization of prophiFSAT01-1, prophiFSIL01-1, McProf, prophiFSQJ01-1 and prophiFVLQ01-1. The ruler represents the coordinates of the genome. Forward and reverse transcribed genes are shown as boxes above and below the ruler, respectively. Genes are colored according to their assigned Phamily with the Phamily number shown above each gene with the number of Phamily members in parentheses. Genome maps were generated using Phamerator and the database, “Actino_Mab_Draft (version 20).” (Cresawn et al. 2011)

